# Barking Up Inferred Trees: Detecting Signatures of Selection in Dog Genomes Using Ancestral Recombination Graphs

**DOI:** 10.64898/2026.09.02.748875

**Authors:** Jeffrey M. Kidd

## Abstract

Identifying the molecular targets of selection is a major goal of evolutionary studies. Prior studies have employed summary statistics to identify genetic signals associated with the domestication of dogs (*Canis lupus familiaris*) from wolves. Recent computational advances enable new approaches based on Ancestral Recombination Graphs, which model the genealogical relationships of samples throughout the genome. Using Ancestral Recombination Graphs, we identified 285 outlier loci with an unusually recent mean time to the most recent common ancestor (TMRCA) in a geographically diverse set of 40 village dogs as compared to 41 wolves. Loci identified from prior studies are enriched in the tails of the observed TMRCA ratio distribution, indicating convergence among diverse approaches for identifying signals of selection in dogs. The outlier loci we identify are within 50 kb of many genes previously linked to domestication and include genes with functions in neurodevelopment and behavior. This study offers a refined set of candidate regions for future functional dissection of the precise molecular changes selected in the early evolution of dogs.

## Introduction

For eons people have wondered how dogs arose from wolves. Analysis of modern and ancient samples shows that dogs (*Canis lupus familiaris*) were domesticated from a now-extinct lineage of wolves 20,000 to 30,000 years ago somewhere in Eurasia [1–3]. Ancient DNA data shows that the diversification of dogs into lineages began at least 14,000 years ago, with at least five major ancestry lineages present by 11,000 years ago [3, 4].

Dissecting the precise genetic differences selected for during domestication has remained a challenge [5, 6]. Animal domestication has been conceptualized as an evolutionary process that may proceed along several pathways with selective pressures that may vary through time [7–9]. Although the degree to which humans directed the earliest stages of domestication is unclear, the process led to dramatic behavioral differences between dogs and wolves, particularly involving their interactions with humans [10–14]. Multiple prior studies have searched for signals in the genome of dogs to identify regions that evolved under selection during domestication, often nominating genes linked to neural function and development, behavior, morphology, diet, and metabolism [15–21].

Since domestication, dogs have experienced a complex demographic history including waves of global migration that parallel the movement of human populations [3]. These demographic events include bottlenecks, expansions, and waves of interbreeding and population replacement that have left distinct signatures in the patterns of canine genetic diversity [22–24]. The formation of modern breeds, largely in the 19^th^ century, had a profound effect on patterns of dog genetic variation [25, 26]. Breed formation also obscured the signals of selection associated with earlier stages of dog evolution [2, 19]. Although these limitations can be partially overcome through the analysis of diverse breed types, comparisons between village dogs and wolves represent an alternative approach. Village dogs are free-ranging populations of dogs found throughout the world that contain the most genetic diversity and represent distinct indigenous populations [27–29]. Scans that compare geographically diverse village dog and wolf populations can therefore detect signatures of selection while limiting the effects of breed formation or local adaptation [2, 19].

Previous studies relied on summary statistics calculated from genetic variation data to identify genomic regions with evidence of non-neutral evolution in dogs [15–21]. However, these statistics offer a noisy summary of the coalescent history of the studied samples. An alternative approach is to base analysis on the Ancestral Recombination Graph (ARG), which represents the full history of coalescence and recombination in a sample of chromosomes [30]. Although reconstructing an ARG is computationally demanding, the past few years have seen major advances in ARG inference, including methods that consider uncertainty in the underlying inferred topologies [31–34]. Many population-genetic parameters of interest can be calculated directly from the branches of the inferred tree, often with less noise than corresponding computations tabulated directly from genotype data [35, 36].

In this study we apply SINGER, a recently described method that samples inferred ARGs, to a collection of village dogs and wolves previously sequenced by the Dog10K Consortium [34, 37]. Using the inferred ARGs, we identify genomic loci with an unusually rapid coalescence, measured as a low mean time to the most recent common ancestor (TMRCA), in village dogs relative to wolves. The identified regions cluster into 285 candidate domestication loci that may have been targets of selection during early stages of dog evolution. The 285 candidate loci are near many genes linked to neural development and that have been previously associated with human neurological and neurodevelopmental disorders.

## Results

### Village dog and wolf population structure

To enrich for signals of selection during early canine domestication, we analyzed a geographically diverse collection of 40 village dogs and 41 wolves previously sequenced by the Dog10K Consortium (Supplementary Table S1). A principal component analysis (PCA) of SNP genotypes cleanly separates dog and wolf samples along PC1, with variation along PC2 correlating with Asian origins in both groups (Supplementary Figure S1).

In a PCA of only the village dogs, the 40 samples from 17 different countries fall into four broad groups (Figure 1A). The first PC separates samples with an East or Southeast Asian origin from the others and includes dogs from Cambodia, China, Myanmar, and Thailand. The remaining samples are separated along PC2. At the top is a collection of samples from Belize, Costa Rica, Fiji, French Polynesia, Mexico, and Peru. Since a previous analysis identified shared European-derived ancestry in dogs from the Americas and the South Pacific, we refer to this group as the European-associated population [28]. The second cluster along PC2 includes dogs from Afghanistan, Azerbaijan, Uzbekistan, and Iran, which we label as Central Asia. Finally, dogs from Africa (Congo, Kenya, and Liberia) extend along PC2, with two samples from Congo being closer to the European-associated group. Although cross-validation shows that K=1 component best fits the data (Supplementary Figure S2), an ADMIXTURE analysis with K=4 components recapitulates these classifications, with a handful of dogs inferred to have ancestry from more than one modeled population (Figure 1B).

**Figure 1.**
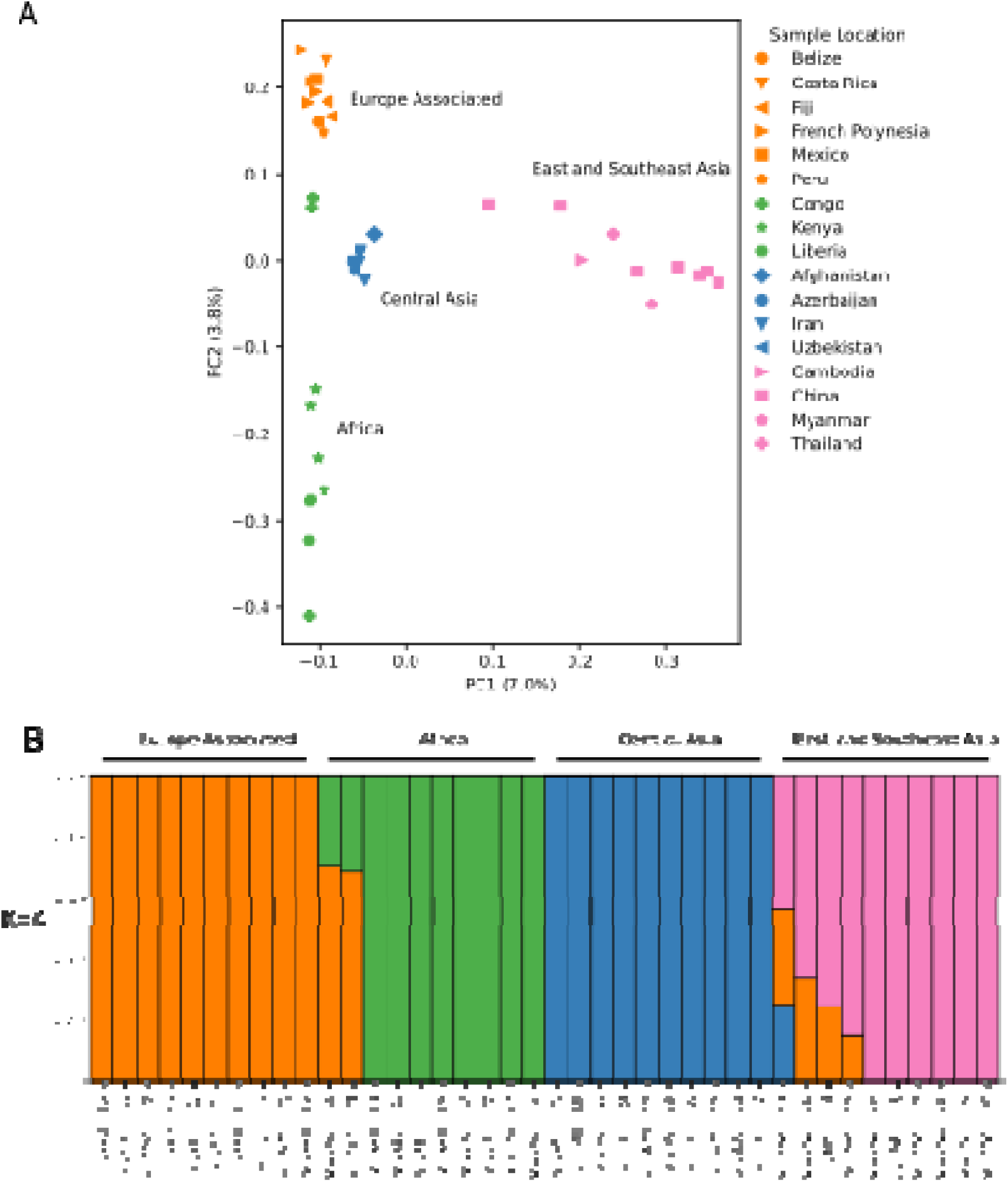
Village dog population structure analysis. (A) The projection of 40 village dogs onto the first two principal components calculated from SNP genotype data is shown. The percent of the total variance explained by each component is given. Samples are colored based on their geographic region. The symbol shapes correspond to countries within each region as indicated. (B) Ancestry results of unsupervised clustering analysis are depicted. Each bar corresponds to one village dog. The colors indicate the proportion of ancestry from each of K=4 modeled source populations.

The available geographic information and apparent clustering for the 41 wolf samples collected from 11 countries is more limited (Supplementary Figure S3). PC1 broadly represents east-west geography, including wolves from Portugal, Greece, Sweden, and eastern portions of Russia (Novosokolnichesky, Tver, and Vesyolovsky) at one end and wolves from northwestern China (Qinghai) at the other. The middle of PC1 features wolves from Tajikistan, Kazakhstan, and northwestern China (Xinjiang). PC2 separates the European cluster from wolves found in Azerbaijan and Iran.

### Candidate selected regions based on TMRCA profiles

Since the above analysis suggests that the samples include a geographically diverse collection of ancestries, we reasoned that comparison between village dogs and wolves may identify loci that evolved under selection during early portions of dog history, prior to the separation of the village dog groups. Using SINGER, we inferred ARGs for village dogs and wolves separately and, for each, calculated the mean pairwise time to the most recent common ancestor (TMRCA) in 1 kb windows [34]. Consistent with a bottleneck shared among dogs, the mean TMRCA for wolves (159,000 generations) is 22% larger than that found in village dogs (130,000 generations) (Figure 2A). The wolf and village dog TMRCAs are strongly correlated along the genome (Pearson r=0.814, p < 0.0001), reflecting their shared history prior to the separation of the ancestral dog lineage (Figure 2B). Following Deng *et al.*, we searched for loci that show differences in coalescence times in wolves and dogs as a signal of selection specific to one population [34]. Specifically, for each 1 kb window we calculated the ratio of the mean TMRCA in wolves to the mean TMRCA found in dogs, taking the mean TMRCA values found across 100 sampled ARGs. Regions where the coalescence in dogs is more rapid than in wolves, leading to a recent village dog TMRCA and an older wolf TMRCA, will have a larger ratio. We identified 2,229 windows in the 0.1% tail of this distribution (corresponding to 39.6 median absolute deviations above the distribution median) as candidates that evolved under selection in dogs (Figure 2C). Although these windows were selected based on the ratio of mean TMRCAs, they correspond to genomic loci with an unusually low TMRCA in village dogs rather than a high TMRCA in wolves (Figure 2D and E). Additionally, a higher fraction of the outlier windows overlaps regions of the genome in which SNPs can be called than expected by chance (98.5% of bp are callable, compared to a 95.5%-97% found in 100 permutations), indicating that false negative variant calls are unlikely to contribute to the observed pattern. We clustered the 2,229 outlier windows into 285 candidate domestication regions spanning a total of 3,073,000 bp (Figure 3, Supplementary Table S2).

**Figure 2.**
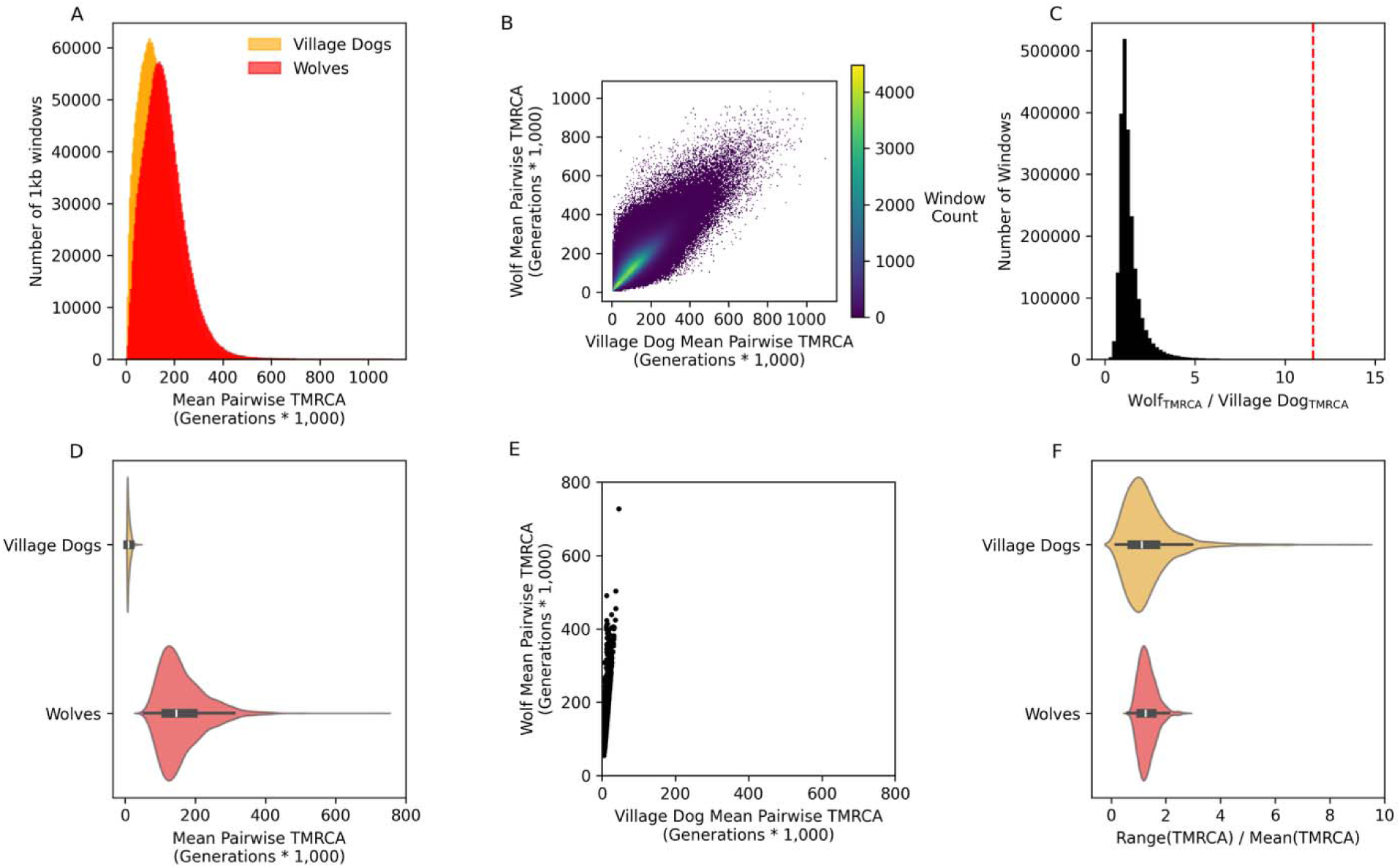
Distribution of mean TMRCA in village dogs and wolves. (A) Histograms of the mean TMRCA for the 40 village dog (orange) and 41 wolf (red) samples calculated in 1 kb windows are shown. (B) A density plot compares the village dog (x-axis) and wolf (y-axis) mean TMRCA values in 1 kb windows through the autosomes. (C) A histogram depicts the distribution of the wolf to village dog TMRCA ratio calculated in 1 kb windows. The red dashed line corresponds to the 0.1% tail. (D) Violin plots show the mean TMRCAs in village dogs (orange) and wolves (red) for the 2,229 1 kb windows with a TMRCA ratio above the cutoff. (E) A scatter plot is shown comparing the mean TMRCA in village dogs (x-axis) and wolves (y-axis) for the 2,229 outlier windows. (F) Violin plots illustrate the variability in TMRCA estimates obtained from 100 sampled ARGs for the 2,229 windows in village dogs and wolves. Variability was calculated as the range of TMRCA estimated for each window across inferred ARGs divided by the mean estimate.

**Figure 3.**
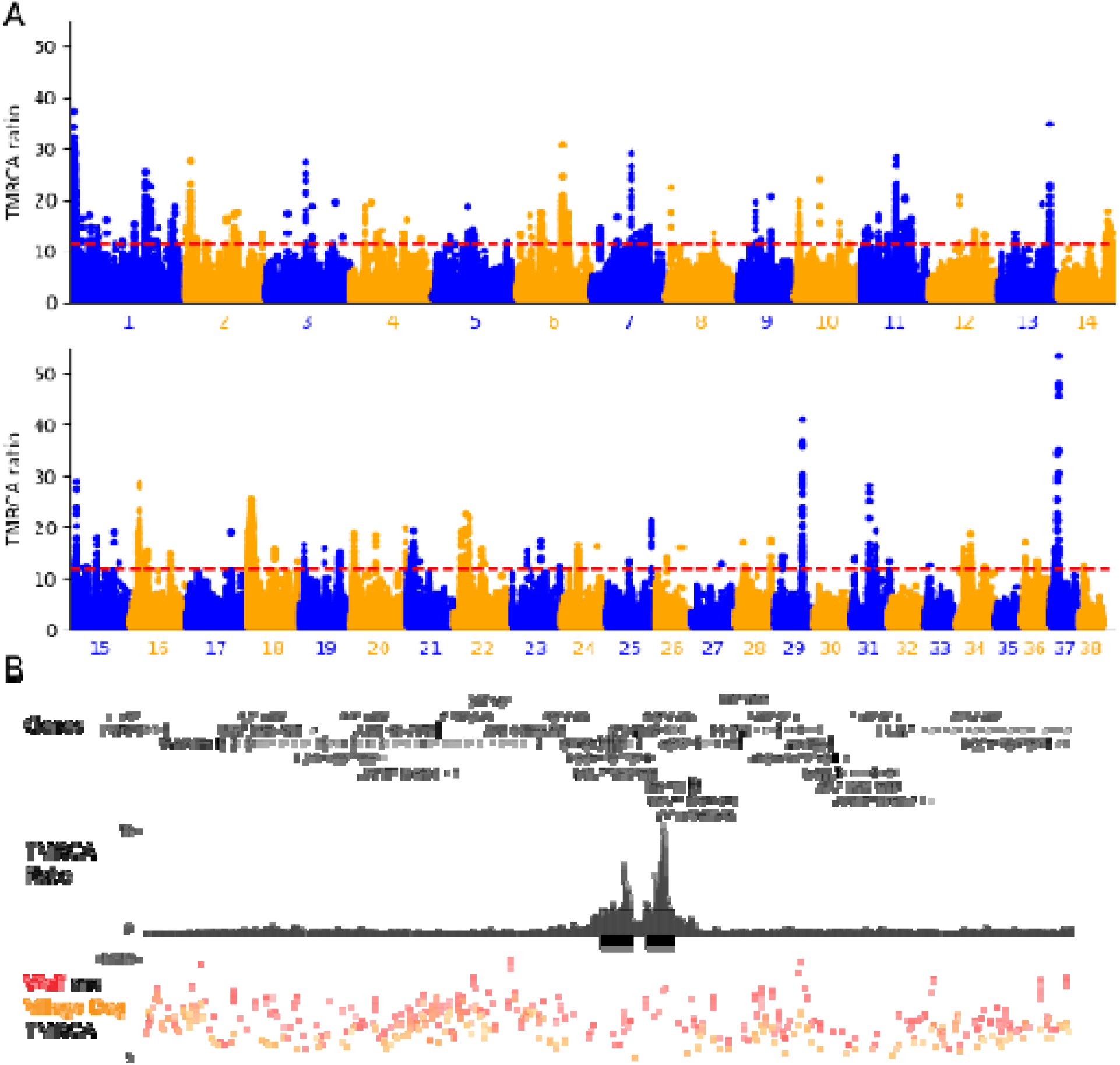
Signature of selection in dog genomes. (A) The ratio of wolf to village dog mean TMRCA plotted in 1 kb windows across the 38 canine autosomes is shown. Chromosomes are depicted in alternating colors as indicated. The dashed red line corresponds to the 0.1% cutoff. (B) A UCSC Genome Browser plot shows a 715 kb region on chr37 that contains the largest detected TMRCA ratio. The top track depicts the position of annotated genes in the region. For each gene, only the longest transcript is shown. The second track shows the Wolf to Village Dog TMRCA ratio. The black horizontal line corresponds to the 0.1% cutoff. The two black rectangles correspond to the candidate domestication regions identified by this study. The bottom track depicts the average mean TMRCA of wolves (red) and village dogs (orange) in generations.

Similar variability in the mean TMRCA estimated from 100 sampled ARGs was found in village dogs and wolves, with the range of estimates for the outlier windows averaging approximately 1.3 times the mean (Figure 2F). To assess the effect of variability in inferred trees, we constructed a conservative ratio using the 25^th^ percentile estimate of the wolf TMRCA and the 75^th^ percentile estimate of the village dog TMRCA found across the 100 sampled ARGs for each of the 2,229 empirical outlier windows. Using this biased estimate, 911 of 2,229 windows exceed the 0.1% empirical cutoff and 2,228 of 2,229 windows exceed the top 1%. The detected windows also have a low mean TMRCA within the four individual village dog geographic groups (Supplementary Figure S4). Across the four groups, 71%-96% of the windows exceed a group-specific 0.1% empirical cutoff with >99% of windows falling within the group-specific 1% tail (Table S3).

### Comparison with previous scans for selection in dogs

We compared the TMRCA ratios we identified with candidate regions identified by previous studies [15–19, 38]. The regions identified in prior studies are enriched in the tails of the TMRCA ratio distribution (Figure 4, Supplementary Table S4). The number and length of loci reported by previous studies varies greatly, with the fraction of loci with a TMRCA ratio exceeding the 0.1% tail correlating with the mean size of the candidates reported by each study (Pearson r=0.917, p=0.01). We therefore assessed the significance of the observed enrichment by randomly permuting the intervals from each study along the genome, finding that the observed enrichments exceed those expected by chance. For example, of the 246 loci (mean length 45.5 kb) identified as candidate domestication regions in our previous study based on the XP-CLR statistic, 36 (14.6%) intersect with a window that exceeds the 0.1% threshold and 82% contain a window with a TMRCA ratio above the 5% tail [19]. This is substantially greater than the 0.7% and 24.2% of intervals expected to fall above each cutoff by chance (Supplementary Table S4).

**Figure 4.**
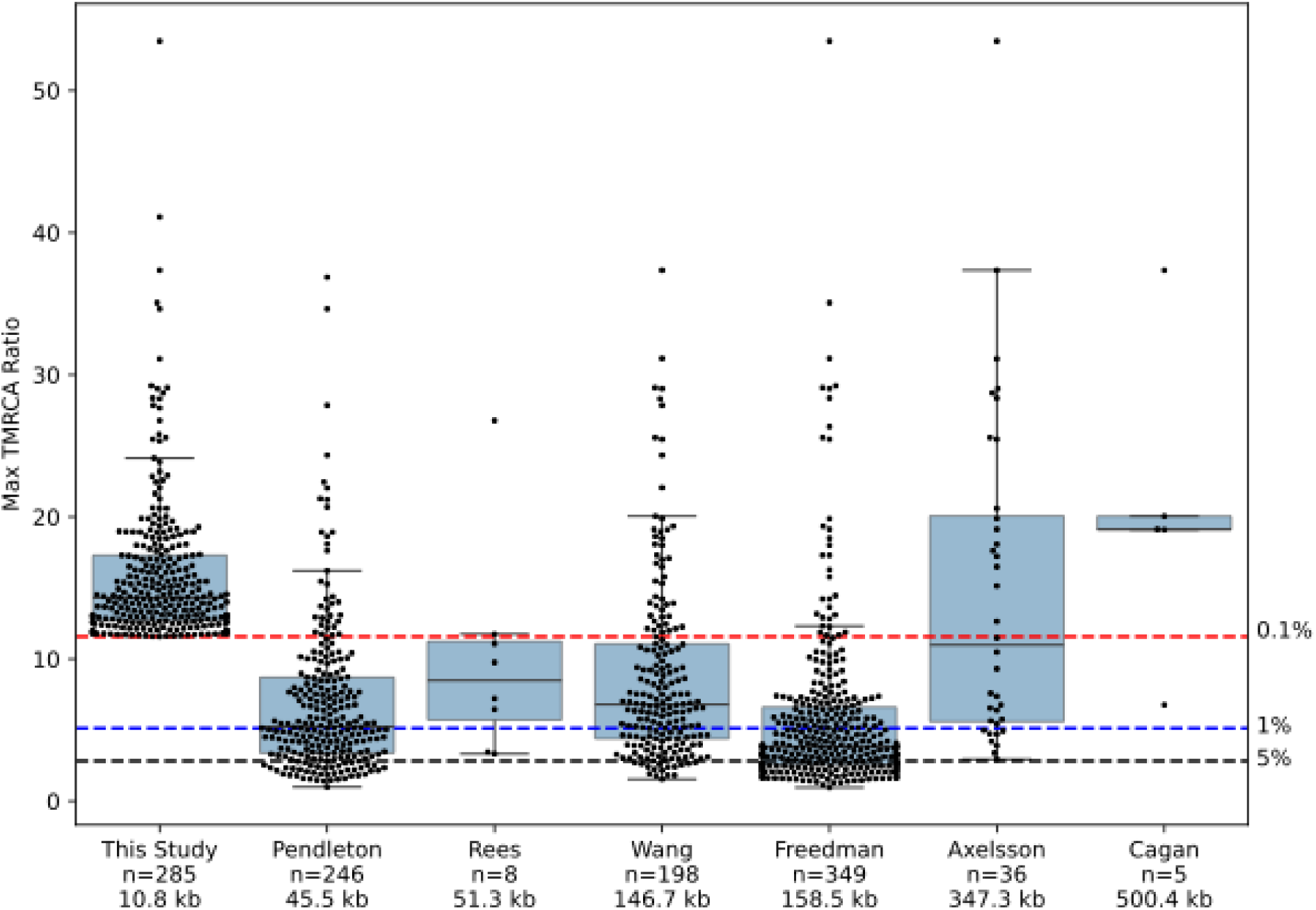
Selected loci identified in previous studies. Swarmplots and boxplots show the maximum TMRCA ratio value found in the selected intervals reported by each study. The dashed lines correspond to the 0.1% (red), 1% (blue), and 5% (black) tails of the observed TMRCA-ratio distribution. The number of loci and the mean locus length reported by each study is indicated. See Supplementary Table S4 for additional information.

Prior studies differed in their design and the type of selection signals they could detect. For example, a recent preprint by Rees *et al.* describes 8 loci identified using a tree-based scan tuned to detect signals of population differentiation between dogs and wolves [38]. This includes a locus with a strong signal in the *MED13L* gene, also identified by Pendleton *et al.* However, this locus is not an outlier in our analysis of mean TMRCAs. Closer examination shows that this region has a reduced mean TMRCA in both village dogs and wolves as well as high F_ST_ between village dogs and wolves, suggesting that different haplotypes have risen in frequency in each group and explaining its absence from our TMRCA-based analysis (Supplementary Figure S5A). In contrast, a region including *IFT88*, which evolved under selection in ancestral wolf populations, shows reduced mean TMRCA in both village dogs and wolves without an elevation in F_ST_ (Supplementary Figure S5B) [39].

### Characteristics of outlier regions identified by TMRCA ratios

The 285 outlier loci we identified are within 50 kb of 396 different genes, which is fewer than expected by chance (mean 658, range 544-849 based on permutation). This set includes multiple neurodevelopmental genes with a putative role in behavior changes. Many of the genes near the outlier intervals are linked to human neurodevelopmental or neuropsychiatric disorders and were also reported by previous studies of dog domestication. For example, the third most extreme TMRCA ratio we detected corresponds to a large region encompassing *GALR1*, which encodes the galanin receptor 1 which is linked to anxiety-like behaviors [40], and *MBP*, which encodes a component of the myelin sheath that surrounds axons [41] (Supplementary Figure S6A). Interestingly, we also identified regions overlapping *MED13*, a member of the Mediator complex, a core transcriptional regulator that is also important for early embryo development and cortical neuron migration (Supplementary Figure S6B) [42–45]. As described above, its paralog, *MED13L*, has reduced TMRCA in both dogs and wolves, suggesting a complex history of selection for key components of transcriptional regulation in canines (Supplementary Figure S5A). Other identified genes are linked to neuronal functions including (***i***) neurotransmitters and receptors (*GAD2, GRIA1*, and *GRIK3* [46–48]), (***ii***) neural development and connectivity (*CADM2*, *NCAM2*, *NRXN3*, and *LRRC4C* [49–52]), and (***iii***) synapse function (*SH3GL2*, *LRRTM4*, *CBLN2*, and *ELFN2* [53–58]).

### Overlap with regions associated with human genomic disorders

Previous studies have identified signals of selection in two regions altered in human genomic disorders. Genomic disorders result from the duplication or deletion of genomic intervals that typically contain multiple genes and are associated with diverse neurodevelopmental phenotypes [59]. The copy-number variable intervals include dosage sensitive genes and serve as examples of regions where regulatory perturbations can give rise to complex phenotypes [60]. Several prior studies have identified *RAI1,* a dosage-sensitive gene that drives the main features of Smith-Magenis and Potocki-Lupski syndromes, as a candidate dog domestication locus [15, 17, 19, 38, 61]. *RAI1* is a chromatin regulator implicated in diverse processes including cell cycle regulation, neural crest cell development and migration, skeletal and craniofacial development, circadian activity, neuronal communication, synaptic plasticity, and behavior [62–66]. Our scan identified two outlier regions within *RAI1* with elevated TMRCA ratios (Supplementary Figure S7A). The regions include predicted dog enhancers [67], further suggesting a role for altered *RAI1* regulation in dog domestication.

A comparison between breed dogs and wolves identified a SNP with high F_ST_ in *GALNT17* (*WBSCR17*), a gene near the Williams-Beuren syndrome critical region [20]. Williams-Beuren syndrome is caused by deletion of a 1.5-1.8 Mb segment containing 26 to 28 genes and is characterized by distinct facial features, cardiovascular and endocrine abnormalities, hypercalcemia, impaired glucose tolerance, developmental delays, increased facial recognition abilities, and hypersociability [68]. Although *GALNT17* is outside of the typical deletion boundaries, the expression levels of flanking genes may be altered in Williams-Beuren syndrome patients [69]. *GALNT17* encodes a N-acetylgalactosaminyl transferase involved in the regulation of cell shape, movement, neurodevelopment, and social behaviors [70, 71]. *GALNT17* may share regulatory interactions with the neighboring gene, *AUTS2*, a gene involved in multiple neurodevelopmental processes [72, 73]. In 2017, a candidate variant analysis in 18 dogs and 10 wolves associated dimorphic mobile element insertions near three genes in this region with human-directed social behaviors [74]. Subsequent analysis of breed dogs with survey-based behavior data identified a polymorphism near *GALNT17* as a predictor of dog social behaviors [75]. *GALNT17* overlaps a region with a high TMRCA ratio and was also reported by a previous study (Supplementary Figure S7B) [17].

### Gene ontology enrichment analysis

We performed a gene ontology enrichment analysis to identify biological pathways that are over-represented among genes within 50 kb of the outlier intervals. No enrichments are significant after correcting for the 17,435 tested pathways. We further explored the functions of genes within 50 kb of the outlier intervals using heuristic filters and identified 16 pathways that (***i***) contain at least three identified genes, (***ii***) have a p-value less than 0.05 calculated using the Parent-Child method, and (***iii***) occur in less than 5% of random permutations (Supplementary Table S5). Regulation of canonical *Wnt* signaling (GO:0060828) is the pathway with the largest number of candidate genes, containing seven genes located near the outlier regions. *Wnt* signaling is a key driver of neural crest cell development [76–78], and disruption of neural crest cell functions may contribute to pleiotropic phenotype changes found in domesticated animals [79]. However, these pathway results should be interpreted with caution.

For example, four genes drive the enrichment of six different GO terms related to the negative regulation of cellular processes (Supplementary Table S5). This set includes a selection signal within an intron of *GLI3*, a gene involved in the Hedgehog signaling pathway that regulates the development of multiple tissues and immune cells [80]. *GLI3* also contributes to the specification and differentiation of neural crest cells and to brain patterning and development [81–84]. In humans, mutations in *GLI3* show pleiotropic effects including polydactyly, craniofacial anomalies, hypothalamic hamartoma, bifid epiglottis, laryngeal cleft, and pulmonary segmentation anomalies [85]. The second signal in this set occurs within a predicted long noncoding RNA (LOC111096980) located between *CD274* and *PDCD1LG2*. These genes encode the PD-L1 and PD-L2 ligands of the PD-1 receptor, a negative regulator of the immune system and a target in cancer immunotherapy [86–88]. The third signal is located in the promoter region of *TMEM131L*, a gene involved in *Wnt* signaling and T-cell development [89]. The fourth signal is located 18 kb downstream of *DUSP3*, a phosphatase linked to diverse functions including immune response, genomic stability, and the maintenance of tight junctions between cells [90, 91]. This signal is also just 5 kb downstream of *SOST*, which makes sclerostin, a protein that inhibits *Wnt* signaling and regulates bone growth [92]. Examination of these six GO categories illustrates how (***i***) uncertainty in linking a selection signal to a specific gene, (***ii***) the pleiotropic nature of gene function, and (***iii***) redundancy among pathway definitions complicate the identification of functional pathways that were targeted by selection.

## Discussion

In this study we used inferred ARGs to identify regions of the genome with TMRCAs suggestive of selection in early stages of dog evolution. To do this, we identified genome intervals where the mean TMRCA among a globally diverse collection of village dogs is substantially lower than that found in wolves. This metric captures differences in coalescence rates at the same locus between populations, a key signal of adaptation, and relies upon the reduced noise of fine-scale diversity metrics represented by ARGs relative to direct computation from variation data [34, 35]. The use of geographically diverse village dogs is designed to enrich for signals from early stages of dog evolution shared across groups and avoids confounding factors associated with the genetic structure of modern breed dogs. Although additional data is required to estimate when selection at each locus began, we refer to these outlier regions as candidate domestication regions in accordance with previous studies and to emphasize our focus on identifying genetic signals from stages of dog evolution prior to the separation of existing lineages.

Multiple previous studies have used a range of statistics and samples to nominate candidate loci that were under selection early in dog evolutionary history. Although there are important differences among studies, we find that the previously reported loci are enriched in the tails of the TMRCA ratio distribution, suggesting that diverse approaches have identified shared signatures to identify loci important for dog evolution.

The outlier regions we identified are near many genes with functions plausibly linked to changes associated with dog domestication, including genes implicated in neurodevelopmental or neuropsychiatric disorders when disrupted in humans. This is relevant not because of putative parallels between phenotypes in humans and other species, but because the study of naturally occurring human genetic variation represents a powerful approach to assign functions to genes and key developmental pathways are broadly shared across mammals. Pathway analysis of the outlier loci is challenging, since existing pathway enrichment approaches are based on the context of gene expression studies and assume that genes are independent and equally likely to be chosen [93]. These assumptions do not hold when analyzing gene sets that intersect with identified genomic regions since genes with similar functions may be clustered together, genes vary dramatically in size, and the largest genes tend to be expressed in neurons [94–96]. Coupled with multiple-hypothesis testing concerns and the non-independence of defined functional pathways, pathway enrichment results should be viewed with caution. Nonetheless, we identified regulation of *Wnt* signaling as the largest over-represented pathway. Although disruption of multiple processes undoubtedly contributed to the evolution of dogs, and each domesticated species has its own history and no one factor can account for all the traits found in domesticates, this finding is consistent with the idea that disruption of neural crest cells contributed to dog domestication [79, 97–99].

There are several potential technical and conceptual limitations to this study. We annotated genes located within 50 kb of the outlier regions, even though regulatory elements can act over much longer distances and a single element may regulate multiple genes [100]. Linking loci to genes and functions remains a major challenge. We also limited our analysis to the autosomes, meaning that we are blind to signals on the X chromosome. This bias may be important since X-linked genes are enriched for expression in the brain [101].

Our analysis is based on SNPs detected by mapping short reads to the UU_Cfam_GSD_1.0/canFam4 assembly derived from a German Shepherd Dog [37, 102]. Although estimates of genetic diversity may be biased when reads are aligned to diverged reference genomes, our previous analysis suggests this is unlikely to affect SNP genotypes obtained from dogs and wolves [103, 104]. Although the inferred ARGs are based on SNP data, they capture variation in coalescence times along the genome that could be due to selection on other variant types. Mobile elements are large contributors to genetic diversity in canines, and dogs may be a valuable test case for the incorporation of mobile elements and other structural variants into ARG-based selection scans [105].

Canine genomes contain duplicated sequences, resulting in segments where genetic variation cannot be confidently identified using short-read data [106]. Some inference programs utilize a mask file to distinguish regions without variation from regions with missing data. Since this option is currently not supported by SINGER, regions in which SNPs cannot be called may be inferred to have a low TMRCA. Because our analysis is based on TMRCA ratios, this is expected to have a minimal impact as long as the rate of missingness is comparable between village dogs and wolves. Although the detected outlier windows are not enriched for regions the Dog10K Consortium identified as problematic for SNP calling, this callable genome mask was based only on Illumina read depth and mapping profiles. More sophisticated metrics that consider variation found in high-quality reference genomes may yield better metrics of the portion of the dog genome from which reliable SNP calls are obtained from short-read data [107].

Our approach identified outliers where a geographically diverse collection of village dogs has an unusually recent TMRCA relative to wolves. Outlier approaches have well-known limitations, and selection scans benefit from models that incorporate demographic history and purifying selection [108, 109]. As ancient DNA studies continue to impose constraints on the domestication process and the early stages of dog diversification it may become possible to develop a robust model of dog demographic history that recapitulates key aspects of dog and wolf genetic variation [3, 39]. This will enable a more sophisticated analysis of the timing and strength of selection acting on individual loci [110].

A major remaining challenge is identifying the specific molecular changes targeted by selection. Previous studies have found a paucity of coding differences between dogs and wolves [19, 21]. The regions we identified are biased away from genes, consistent with a predominant role for selection on gene regulatory differences. Functionally dissecting these changes will be aided by the comparably small size of the candidate loci we identified (mean length of 10.8 kb) and the continued development of dog regulatory maps [67, 111]. However, functional studies should be designed with caution since some candidate loci are characterized by extended regions of low TMRCA in dogs with the boundaries of the reported loci reflecting variation in the wolf TMRCA along the interval. Additionally, it is important to consider wolf gene regulatory features that may have been lost in dogs [112]. An ambitious long-term goal is linking molecular changes to specific genes and understanding the associated changes in gene functions in the context of the organismal traits selected for during dog evolution. In the coming years the combination of comprehensive catalogs of canine genetic variation derived from long reads, which capture both sequence and structural variation, appropriate cellular models, and massively parallel reporter assays is likely to identify precise molecular changes that were selected for during dog domestication [113–115].

## Methods

### Phasing and population structure analysis

Phased and imputed genotypes were determined for SNPs identified in 1,929 dog and wolf samples by the Dog10K Consortium using SHAPEIT5 (v5.1.1) [37, 116]. A version of the fine-scale genetic map inferred by Auton *et al.,* converted to canFam4 coordinates, was used [117, 118]. Due to differences in the evolutionary history of the X chromosome, all analyses were limited to the 38 autosomes. Only biallelic sites were considered, resulting in a total of 29,307,684 SNPs.

From the Dog10K data, we selected a set of 40 village dogs, including ten samples each from four broad geographic categories. Previous analysis suggested that the wolf populations in Greece and Sweden have experienced a stronger recent population decline than others [104]. We therefore selected one sample each from these two countries. Along with the 39 remaining wolves in the Dog10K collection, this resulted in a set of 41 wolf samples. Sample names are given in Supplementary Table S1.

Sites variable in the 40 village dog (12,624,910 SNPs) and 41 wolf (16,385,832 SNPs) samples were extracted using bcftools (v1.21) [119]. Estimates of F_ST_ between village dogs and wolves were calculated in 20 kb windows using vcftools (version 0.1.15) [120]. For population structure analysis, sites with a minor allele frequency below 5% and in strong linkage disequilibrium were removed using plink (v 1.9) (commands --maf 0.05 and --indep-pairwise 50 5 0.2) [121]. Filtering was performed separately for the combined set of 81 samples and for dogs and wolves individually. Principal component analysis (PCA) was performed using the smartpca tool from the EIGENSOFT (v8.0.0) package [122]. A model-based analysis of population structure was performed using ADMIXTURE (v.1.4.0) [123]. Five runs were performed for each K value and the run with the lowest cross-validation error was used for visualization.

### Ancestral Recombination Graph inference

Ancestral Recombination Graphs (ARGs) were inferred using SINGER (v0.1.9) with the Auton *et al.* recombination map converted to canFam4 coordinates [34, 118]. ARGs were inferred separately for village dogs and wolves. We assumed a mutation rate of 4.5*10^-9^ bp/generation estimated from a wolf pedigree [124]. This rate is compatible with estimates calibrated using ancient DNA and is slightly lower than an estimate from breed dog pedigrees [1, 2, 125]. The effective population size parameter used in ARG inference was estimated from the observed average pairwise village dog and wolf diversity calculated from the site frequency spectrum (village dogs: N_e_ 67,000, wolves: N_e_ 85,000) assuming a mutation rate of 4.5*10^-9^ bp/generation and using the total length of the autosomal chromosomes (2,228,550,668 bp). Sites were assumed to be unpolarized with respect to ancestral state (option-polar 0.5). ARGs were inferred across the autosomes in non-overlapping 5 Mb segments. A total of 120 posterior samples were generated for each segment, with 50 MCMC iterations between samples (options -n 120 and -thin 50). The first 20 samples were discarded as a burn-in period, resulting in 100 sampled ARGs retained for analysis. For each chromosome, inferred ARGs from each posterior sample were converted to the tree sequence format and concatenated using tskit (v 1.0.2) [126, 127]. Examination of the deviation in fit to the diversity landscape and the number of sites incompatible with the inferred tree across 6,000 iterations for 40 village dogs shows that SINGER rapidly reaches stationarity for this data set (Supplementary Figure S8).

### Detection of outliers based on TMRCA ratios

Mean pairwise divergence in units of generations was calculated in 1 kb windows using the ts.diversity function (mode=’branch’) and converted to mean TMRCA by dividing by two. For each window, the mean divergence across the 100 sampled ARGs was calculated. The TMRCA ratio was calculated as the mean TMRCA in wolves divided by the mean TMRCA in village dogs. Windows in the 0.1% tail (ratio value of 11.5765) were identified as outlier candidate selected loci. Windows within 10 kb were merged together using bedtools (v2.30.0), resulting in 285 candidate domestication regions [128] (Supplementary Table S2). The identified windows were intersected with the callable genome mask of regions with read-mapping profiles consistent with the ability to call SNPs by the Dog10K Consortium and compared with the overlap found in 100 window permutations using bedtools shuffle (v2.30.0).

To assess robustness we also calculated a conservative ratio for each outlier window by dividing the 25^th^ percentile of the wolf mean TMRCA estimate by the 75^th^ percentile of the village dog mean TMRCA estimate. We additionally calculated mean TMRCA times within each of the four geographically distinct village dog populations (10 samples each), calculated from the existing ARGs inferred for all 40 samples. The resulting TMRCA divergence ratios relative to wolves were compared with empirical cutoffs calculated based on TMRCA ratios for each population.

### Comparison with previous studies

Regions reported to be evolving under selection early in the history of dogs were obtained from six previous studies [15–19, 38]. Coordinates were converted to the canFam4 assembly using the UCSC liftOver tool [129]. Regions that failed liftOver conversion were manually remapped using minimap2 (v 2.26) to construct the most contiguous aligned interval [130, 131]. The converted interval coordinates were intersected with the TMRCA ratio estimates and the largest ratio value observed for each interval was retained. Enrichment of reported intervals for values in the 0.1%, 1% and 5% tail of the observed TMRCA ratio distribution was calculated based on 100 permutations using bedtools shuffle (v2.30.0), with placement restricted to the autosomes. The mean number of intervals with a ratio above the cutoff was calculated. We also determined how many permutations contained at least as many intervals above the cutoff as found in the original data. A permutation p-value was calculated as 1 plus this number divided by 101.

### Gene analysis

Genes within 50 kb of each annotated candidate locus were identified using bedtools (v2.30.0). The NCBI gene annotation (release 106) modified by the Dog10K Consortium to remove duplicate copies was used [37]. The expected number of intersecting genes was determined by randomly permuting the coordinates of the 285 selected loci using bedtools shuffle 1,000 times, limiting analysis to the autosomes.

Gene ontology enrichment was calculated using topGO (v 2.62.0, R version 4.5.1) with the Parent-Child method using GO annotations from the EBI Gene Ontology Annotation Database (goa_dog.gaf, version 2026-05-19) [132–134]. The set of all autosomal genes with a GO annotation was used as the background. The number of selected genes belonging to each GO category was determined for each of 1,000 random placements of the selected intervals. A permutation-based p-value for each GO term was defined as one plus the number of permutations containing at least as many genes as observed in the real data divided by 1,001 [135]. GO terms passing the following heuristics were identified: (***i***) the category contained at least three genes identified from the selection scan, (***ii***) the pathway had a Parent-Child p-value less than 0.05, and (***iii***) the pathway had a permutation p-value less than 0.05.

## Data access

A UCSC browser TrackHub reporting the TMRCA values and selection intervals is available at https://github.com/KiddLab/village-dog-arg-selection. Biallelic SNPs phased from 1,929 samples using SHAPEIT5 along with village dog and wolf TMRCAs in 1 kb windows are available from the Zenodo data repository at https://zenodo.org/records/22014728.

## Acknowledgements

We are grateful to Susanna Gutierrez, Michelle Kim, Emily Koch, Noel McAllister, and Xinjun Zhang for comments and discussion related to this study.

## Tables

### Supplementary Tables

**Supplementary Table S1.** Analyzed samples.

**Supplementary Table S2.** Candidate selected regions.

**Supplementary Table S3.** Outlier windows in each village dog population.

**Supplementary Table S4.** Comparison of regions identified by previous scans for selection

**Supplementary Table S5.** Gene ontology enrichment analysis

## Supplementary Figures

**Supplementary Figure S1.**
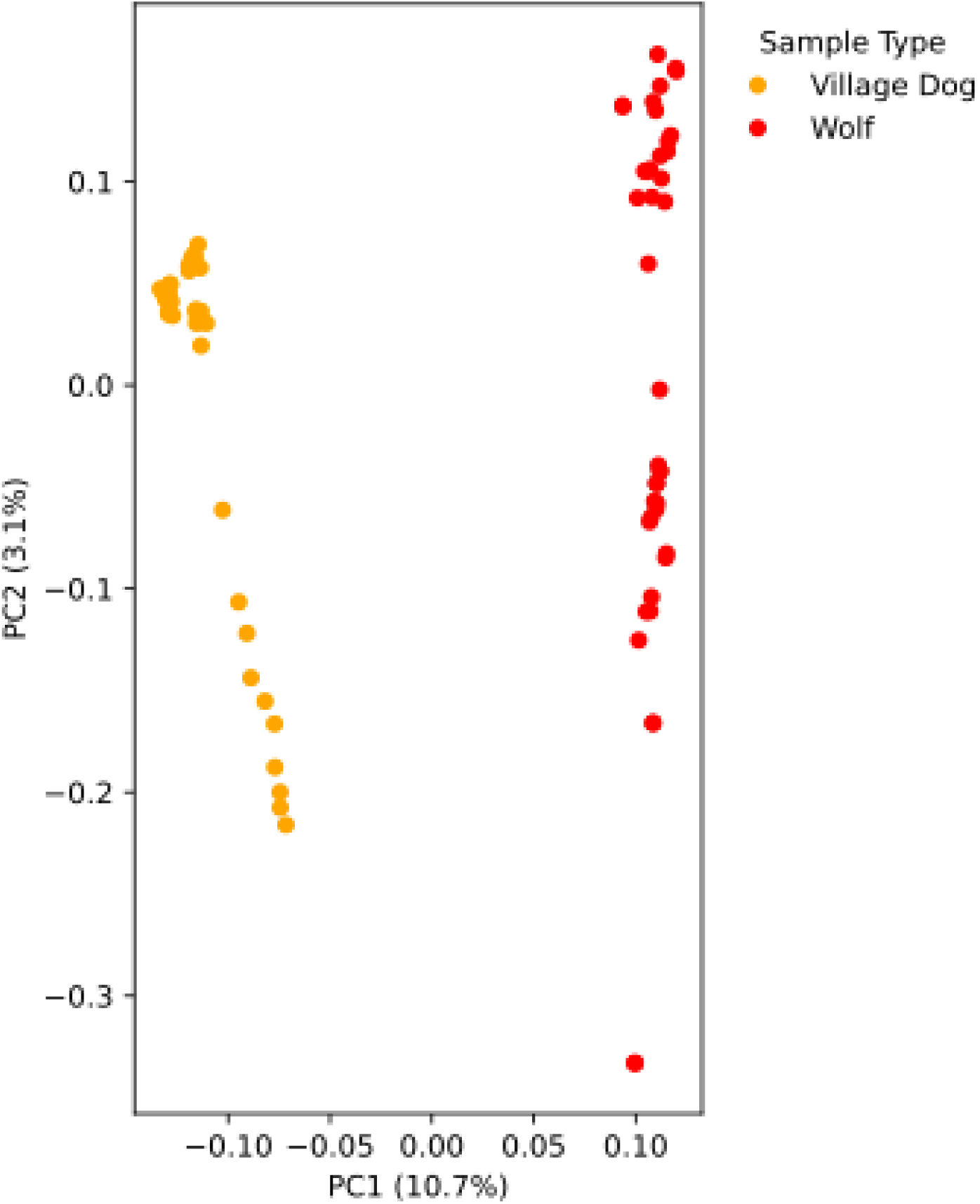
PCA of Village Dogs and Wolves. The projection of 40 village dogs and 41 wolves onto the first two principal components is shown. The percent of the total variance explained by each component is given.

**Supplementary Figure S2.**
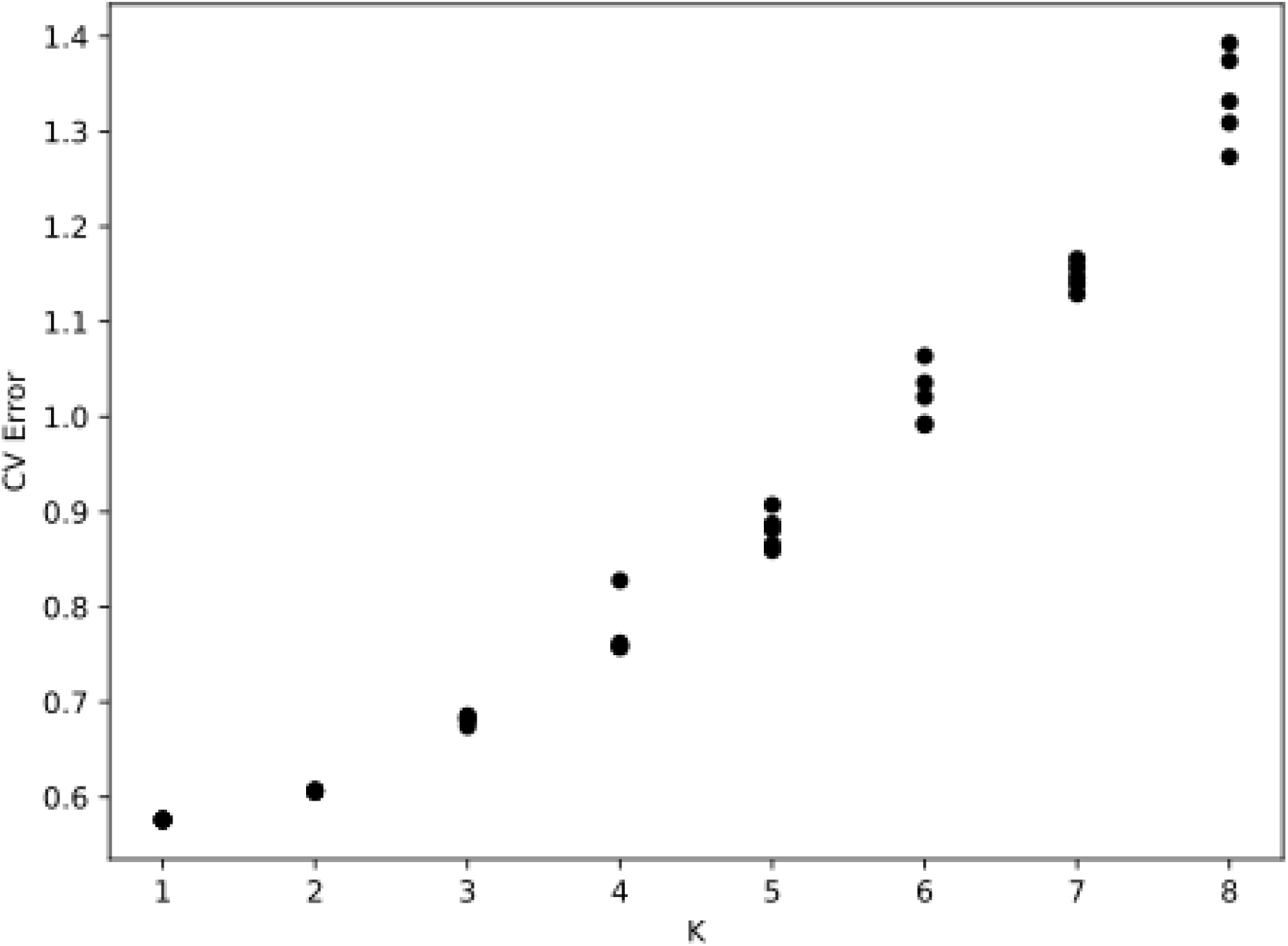
Cross-validation results for village dog ADMIXTURE analysis. Cross-validation errors are shown for analysis of 40 village dog samples assuming K=1 to 8 ancestry components. Five independent runs were performed for each K value.

**Supplementary Figure S3.**
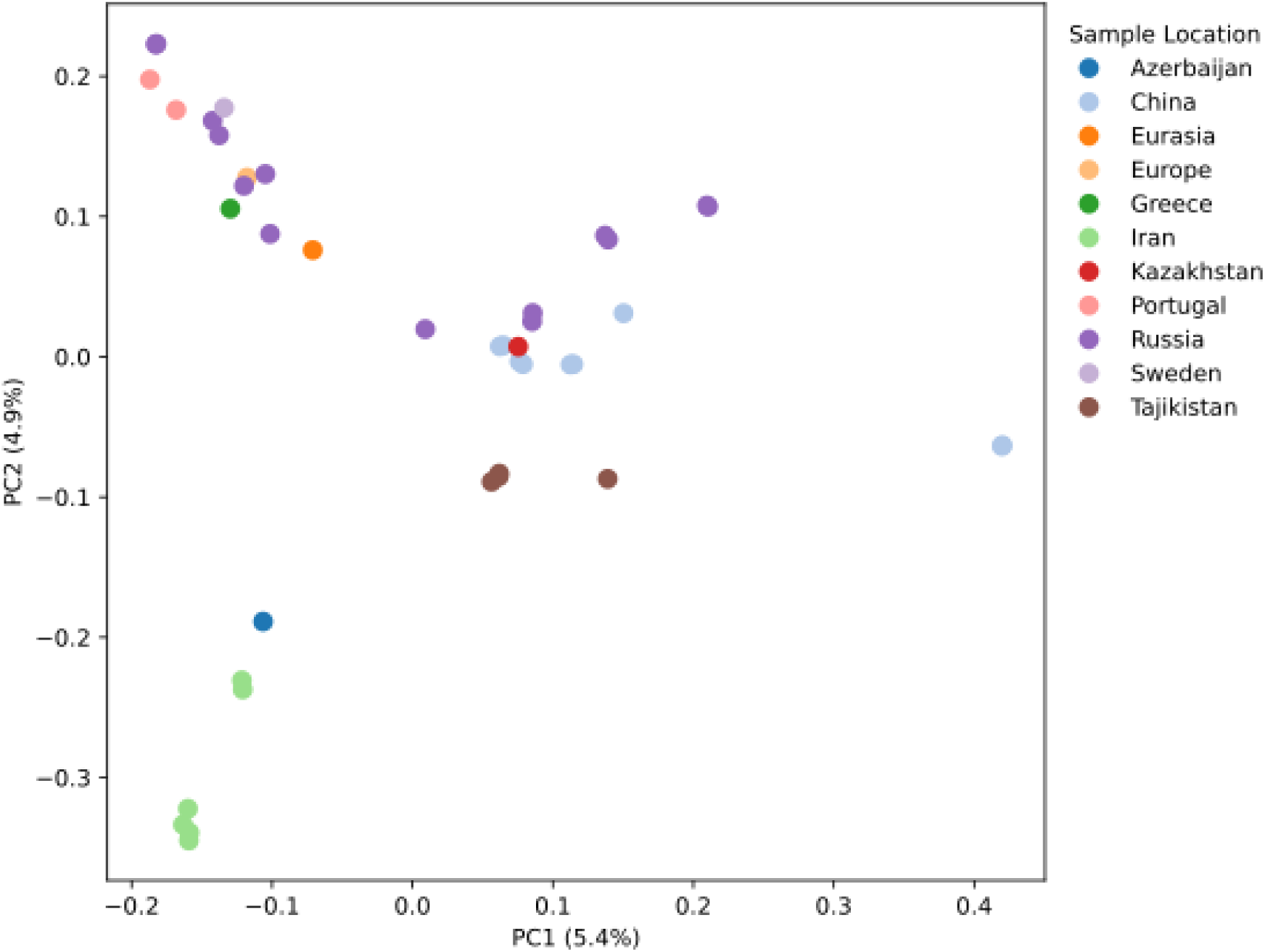
PCA of wolves. The projection of 41 wolves onto the first two principal components is shown. The percent of the total variance explained by each component is given. Samples are colored by the country of origin as indicated.

**Supplementary Figure S4.**
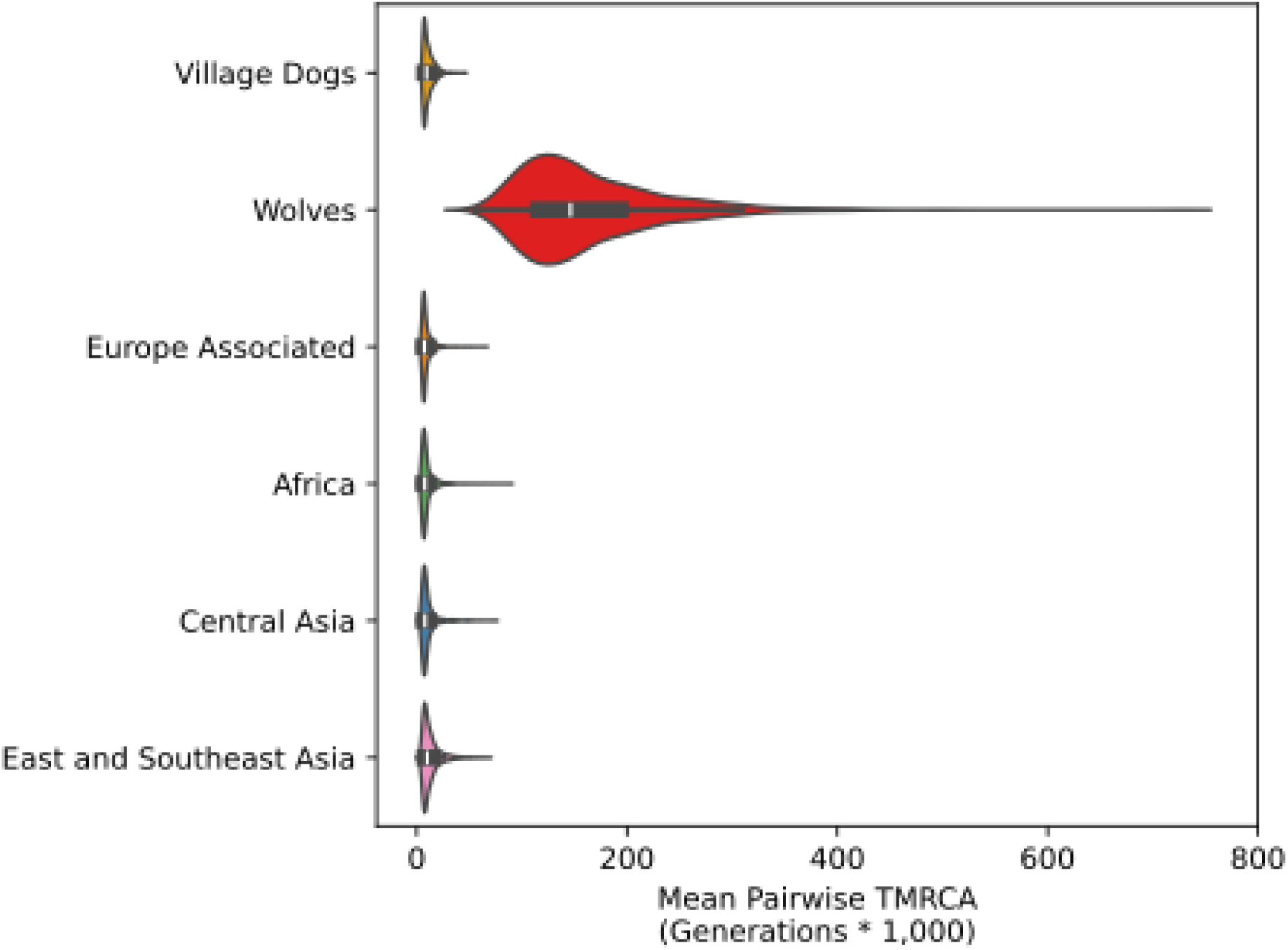
Mean TMRCA within each village dog group. Violin plots depict the mean TMRCA found for the 2,229 outlier windows. Values are shown for 40 village dogs, 41 wolves, and 10 village dogs from each of four globally diverse groups.

**Supplementary Figure S5.**
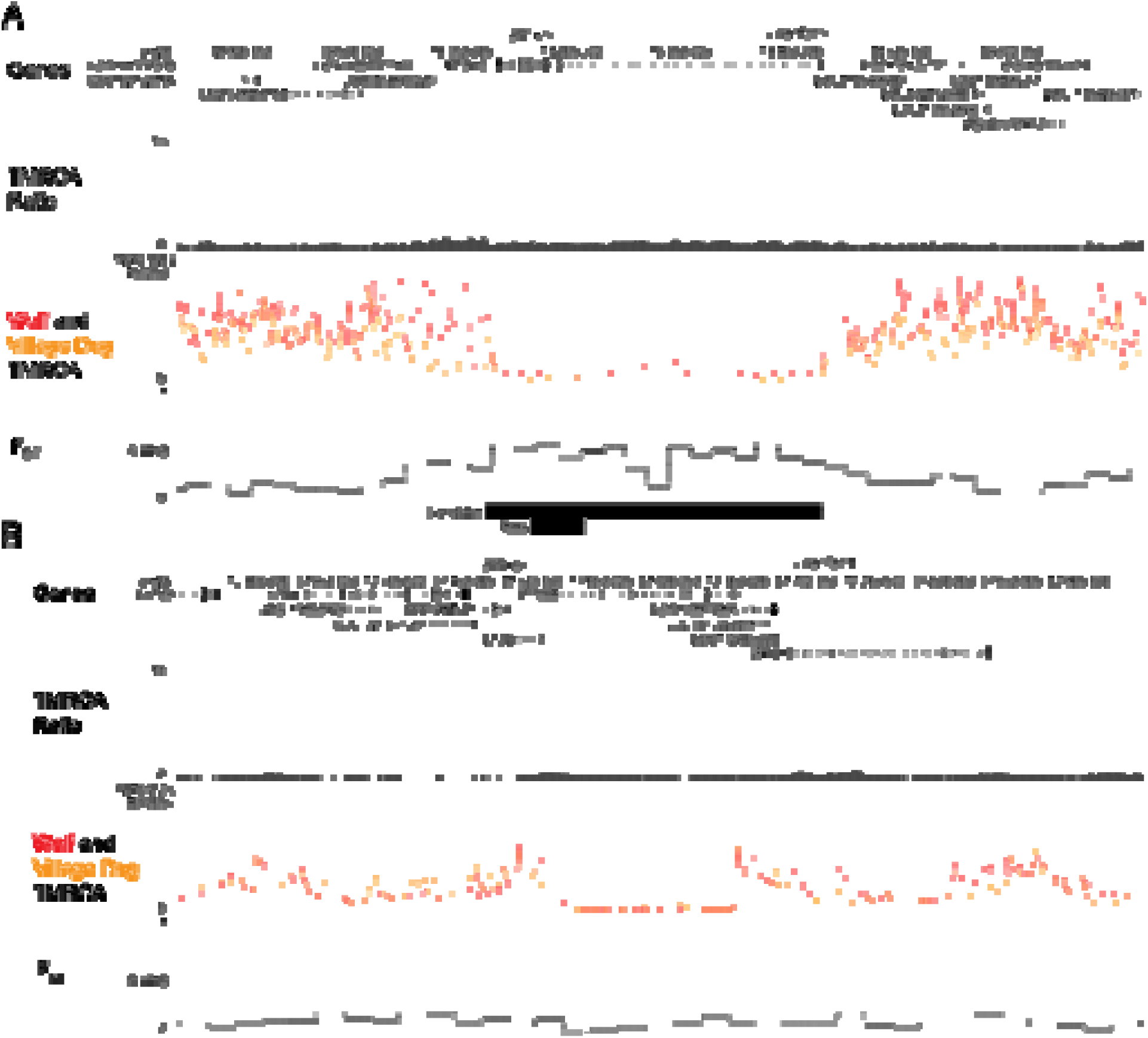
Regions with a signature of selection in both village dogs and wolves. UCSC Genome Browser plots are shown for two regions with reduced TMRCA in both village dogs and wolves. (A) A 875 kb region on chr26 containing *MED13L* is shown. As in Figure 3, the top track depicts the position of annotated genes in the region. For each gene, only the longest transcript is shown. The second track shows the Wolf to Village Dog TMRCA ratio. The black horizontal line corresponds to the 0.1% cutoff. The third track depicts F_ST_ estimated from SNP data in 20 kb windows. The horizontal line depicts the 1% tail of the observed F_ST_ distribution. The two rectangles at the bottom of the plot correspond to candidate selected regions reported by Pendleton *et al.* and Rees *et al.* (B) A 700 kb region on chr25 containing *IFT88* is shown. Tracks are as in panel A.

**Supplementary Figure S6.**
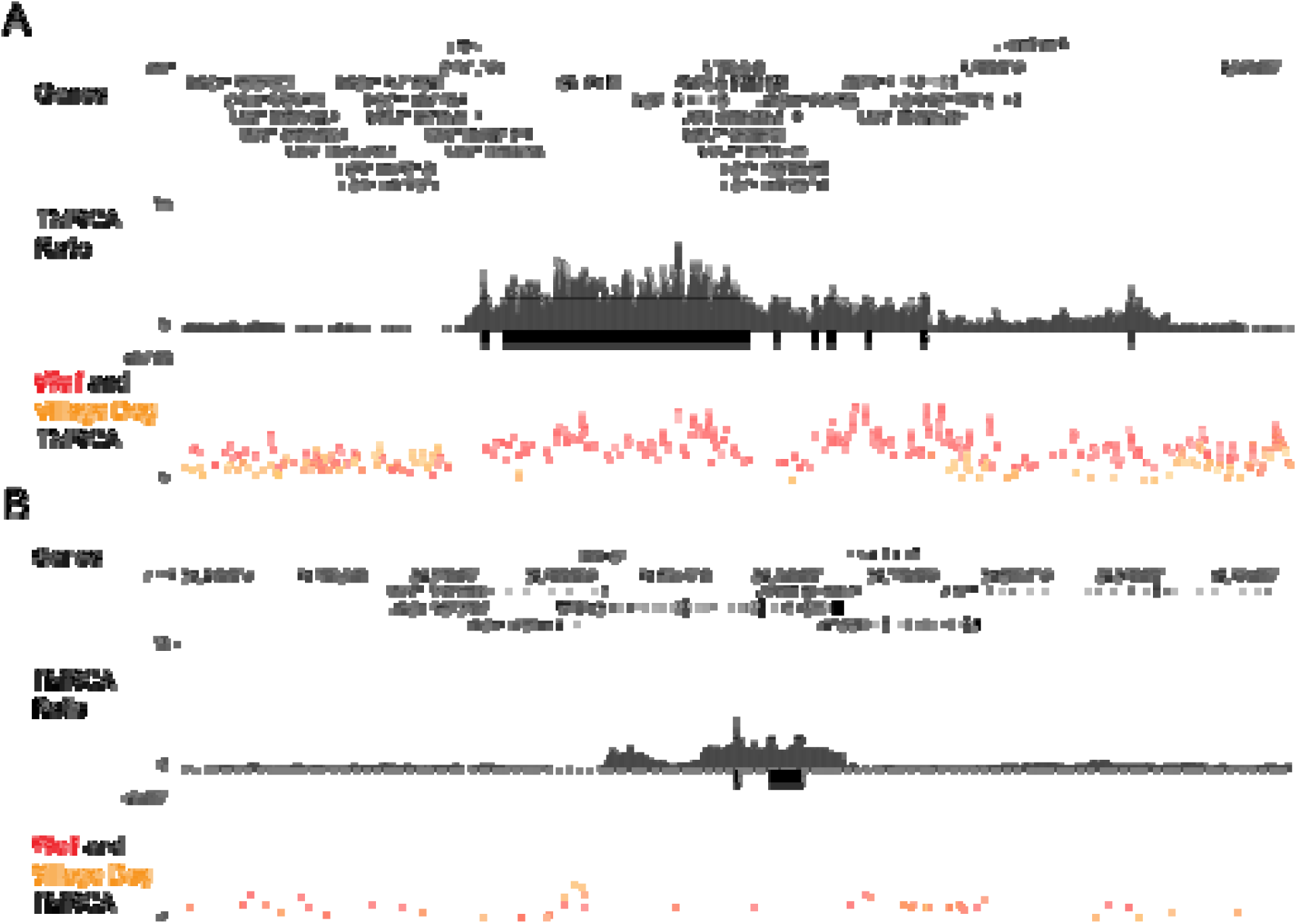
Selection signals at *GALR1, MBP,* and *MED13* UCSC Genome Browser plots are shown for two candidate domestication loci. (A) A 2.1 Mb region on chr1 intersecting with *GALR1* and *MBP* is shown. (B) A 500 kb region on chr9 overlapping with *MED13* is shown. The tracks are as in Figure 3B.

**Supplementary Figure S7.**
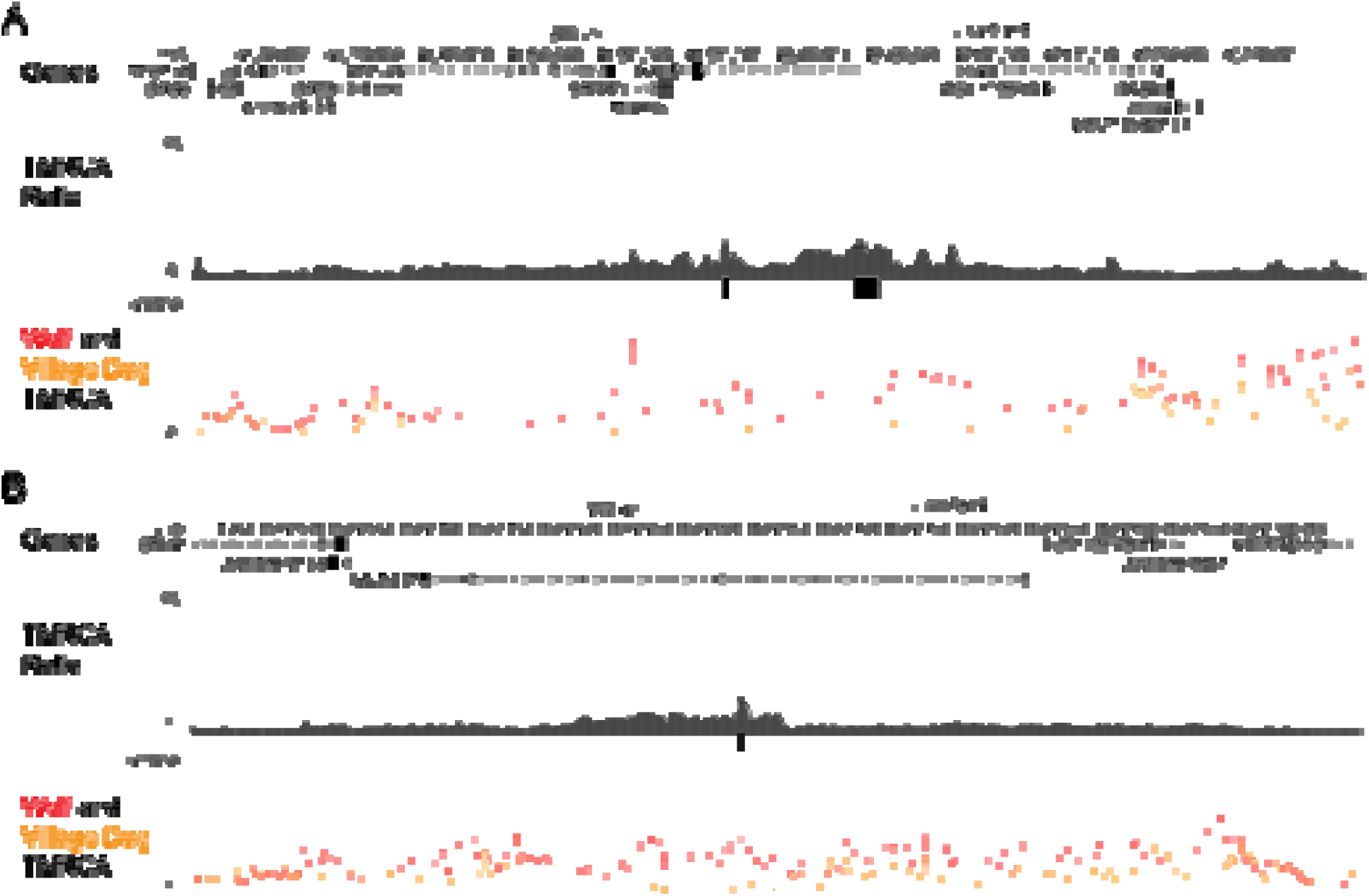
Signals of selection near regions associated with human genomic disorders. UCSC Genome Browser plots are shown for candidate domestication loci in regions linked to genomic disorders. (A) A 650 kb region on chr5 containing *RAI1* is shown. This locus contains two regions identified as candidate domestication regions in this study. *RAI1* is the main gene associated with Smith-Magenis and Potocki-Lupski syndromes. (A) An 850 kb region on chr6 containing *GALNT17* is shown. This locus contains one region identified as a candidate domestication region in this study. The region is adjacent to the critical region associated with Williams-Beuren syndrome in humans. The tracks are as in Figure 3B.

**Supplementary Figure S8.**
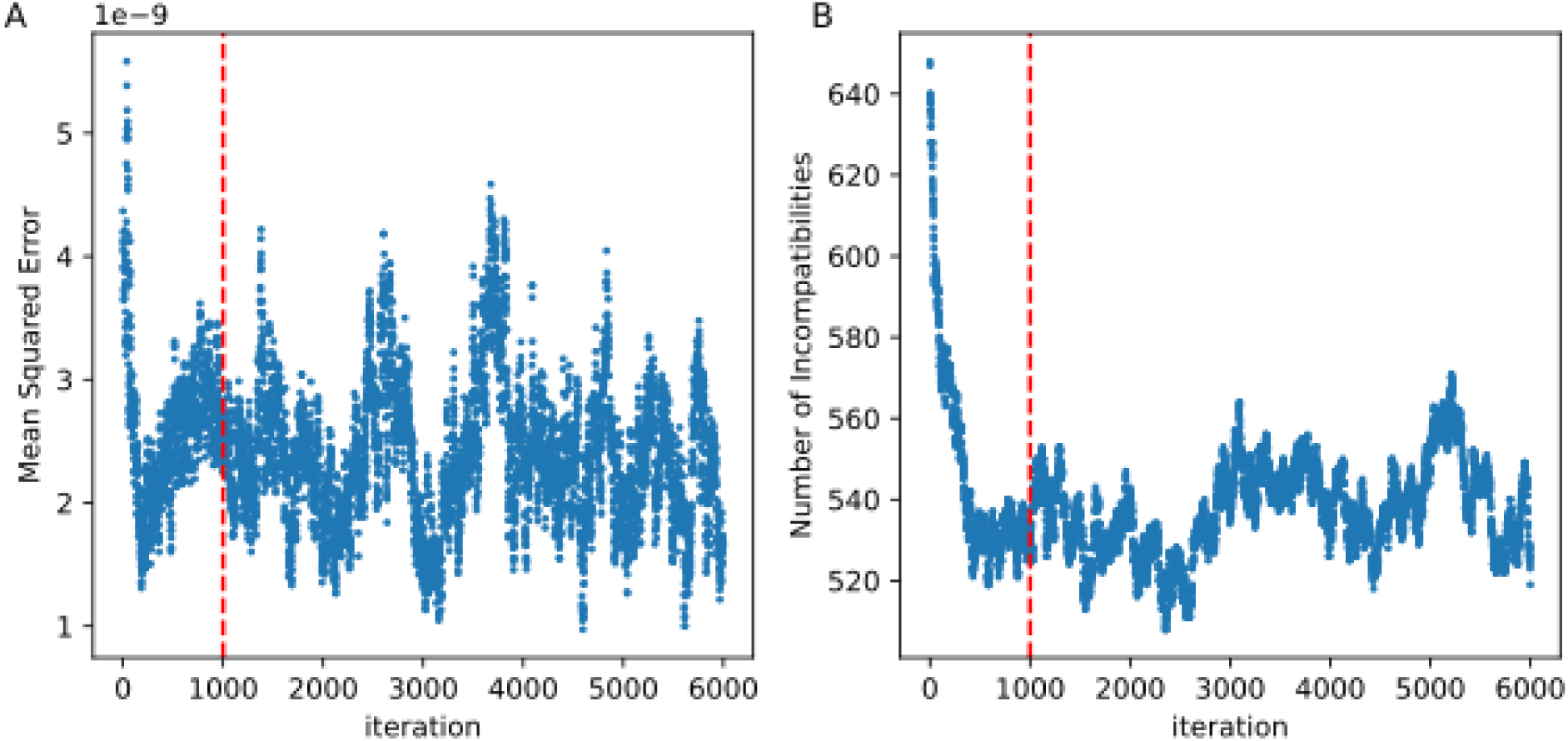
SINGER stationarity trace plots. Trace plots are shown for (A) the mean squared error of deviation in fit to the diversity landscape and (B) the number of incompatible sites for 6,000 iterations. ARGs were inferred for 40 village dogs in the first 5 Mb of chr36. The red dashed line indicates the burn-in period of 1,000 iterations.

