## Supplementary Figures for "Barking Up Inferred Trees: Detecting Signatures of Selection in Dog Genomes Using Ancestral Recombination Graphs"

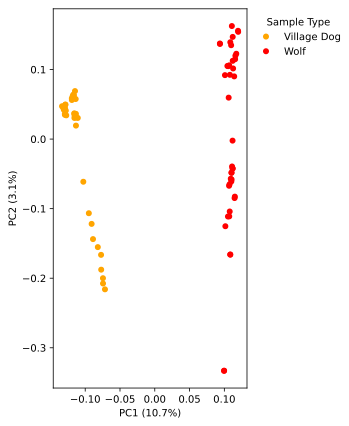

**Supplementary Figure S1.** PCA of Village Dogs and Wolves.

The projection of 40 village dogs and 41 wolves onto the first two principal components is shown. The percent of the total variance explained by each component is given.

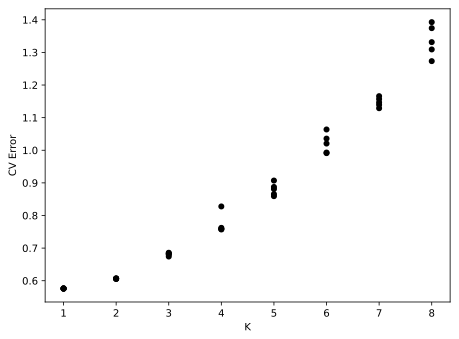
**Supplementary Figure S2.** Cross-validation results for village dog ADMIXTURE analysis.

Cross-validation errors are shown for analysis of 40 village dog samples assuming K=1 to 8 ancestry components. Five independent runs were performed for each K value.

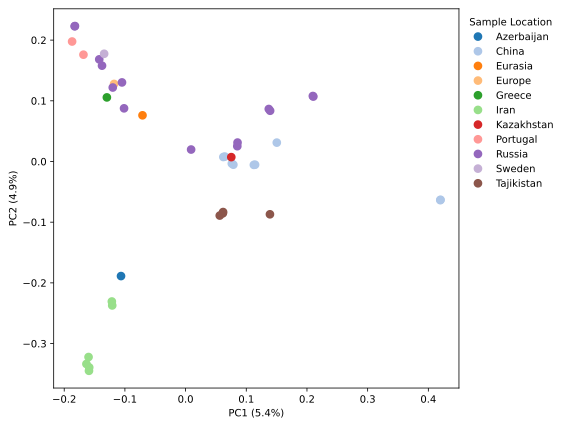

**Supplementary Figure S3.** PCA of wolves.

The projection of 41 wolves onto the first two principal components is shown. The percent of the total variance explained by each component is given. Samples are colored by the country of origin as indicated.

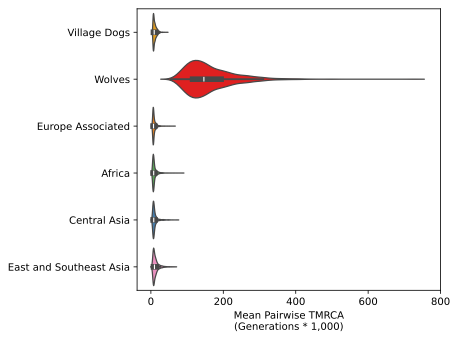

**Supplementary Figure S4.** Mean TMRCA within each village dog group

Violin plots depict the mean TMRCA found for the 2,229 outlier windows. Values are shown for 40 village dogs, 41 wolves, and 10 village dogs from each of four globally diverse groups.

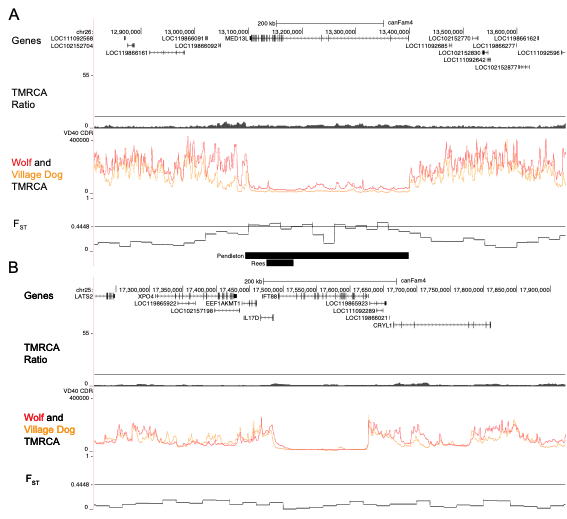

**Supplementary Figure S5.** Regions with a signature of selection in both village dogs and wolves.

UCSC Genome Browser plots are shown for two regions with reduced TMRCA in both village dogs and wolves.

(A) A 875 kb region on chr26 containing *MED13L* is shown. As in Figure 3, the top track depicts the position of annotated genes in the region. For each gene, only the longest transcript is shown. The second track shows the Wolf to Village Dog TMRCA ratio. The black horizontal line corresponds to the 0.1% cutoff. The third track depicts F_ST_ estimated from SNP data in 20 kb windows. The horizontal line depicts the 1% tail of the observed F_ST_ distribution. The two rectangles at the bottom of the plot correspond to candidate selected regions reported by Pendleton *et al.* and Rees *et al.* (B) A 700 kb region on chr25 containing *IFT88* is shown. Tracks are as in panel A.

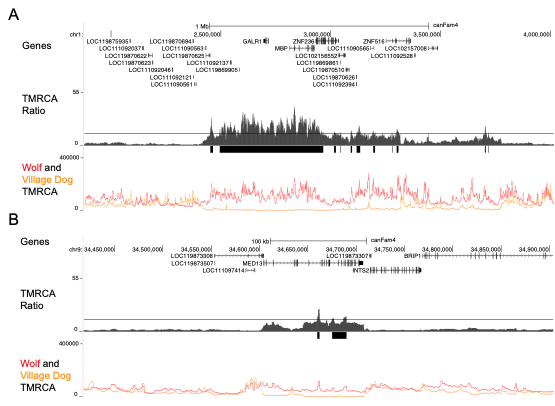

**Supplementary Figure S6.** Selection signals at *GALR1, MBP,* and *MED13*

UCSC Genome Browser plots are shown for two candidate domestication loci. (A) A 2.1 Mb region on chr1 intersecting with *GALR1* and *MBP* is shown. (B) A 500 kb region on chr9 overlapping with *MED13* is shown. The tracks are as in Figure 3B.

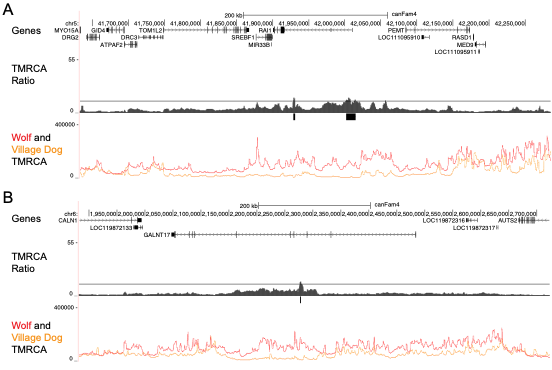

**Supplementary Figure S7.** Signals of selection near regions associated with human genomic disorders.

UCSC Genome Browser plots are shown for candidate domestication loci in regions linked to genomic disorders. (A) A 650 kb region on chr5 containing *RAI1* is shown. This locus contains two regions identified as candidate domestication regions in this study. *RAI1* is the main gene associated with Smith-Magenis and Potocki-Lupski syndromes. (B) An 850 kb region on chr6 containing *GALNT17* is shown. This locus contains one region identified as a candidate domestication region in this study. The region is adjacent to the critical region associated with Williams-Beuren syndrome in humans. The tracks are as in Figure 3B.

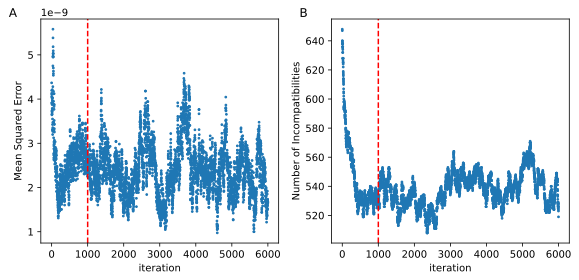

**Supplementary Figure S8.** SINGER stationarity trace plots

Trace plots are shown for (A) the mean squared error of deviation in fit to the diversity landscape and (B) the number of incompatible sites for 6,000 iterations. ARGs were inferred for 40 village dogs in the first 5 Mb of chr36. The red dashed line indicates the burn-in period of 1,000 iterations.
